# Enzyme annotation in UniProtKB using Rhea

**DOI:** 10.1101/709899

**Authors:** Anne Morgat, Thierry Lombardot, Elisabeth Coudert, Kristian Axelsen, Teresa Batista Neto, Sebastien Gehant, Parit Bansal, Jerven Bolleman, Elisabeth Gasteiger, Edouard de Castro, Delphine Baratin, Monica Pozzato, Ioannis Xenarios, Sylvain Poux, Nicole Redaschi, Alan Bridge, The UniProt Consortium

## Abstract

**Motivation:** To provide high quality computationally tractable enzyme annotation in UniProtKB using Rhea, a comprehensive expert-curated knowledgebase of biochemical reactions which describes reaction participants using the ontology ChEBI (Chemical Entities of Biological Interest).

**Results:** We replaced existing textual descriptions of biochemical reactions in UniProtKB with their equivalents from Rhea, which is now the standard for annotation of enzymatic reactions in UniProtKB. We developed improved search and query facilities for the UniProt website, REST API, and SPARQL endpoint that leverage the chemical structure data, nomenclature, and classification that Rhea and ChEBI provide.

**Availability and Implementation:** UniProtKB at https://www.uniprot.org/; UniProt REST API at https://www.uniprot.org/help/api; UniProt SPARQL endpoint at https://sparql.uniprot.org/sparql; Rhea at https://www.rhea-db.org/.

## 1 Introduction

The UniProt Knowledgebase (UniProtKB, at https://www.uniprot.org) is a reference resource of protein sequences and functional annotation that covers proteins from all branches of the tree of life (The UniProt Consortium, 2019). UniProtKB includes an expert curated core of over 560,000 reviewed UniProtKB/Swiss-Prot protein sequence entries that is supplemented by over 158 million unreviewed UniProtKB/TrEMBL entries annotated by automatic systems (release 2019_05 of June 5 2019). UniProtKB/Swiss-Prot curation focuses on experimentally characterized proteins from a broad range of taxa, including proteins of human origin (Breuza, et al., 2016) as well as proteins of bacteria, archaea, viruses, and plants. Approximately half of all protein sequence entries in UniProtKB/Swiss-Prot describe enzymes, whose function has traditionally been annotated using reference vocabularies such as the hierarchical enzyme classification of the Enzyme Nomenclature Committee of the IUBMB (often referred to as EC numbers) (Bairoch, 2000; McDonald, et al., 2009; McDonald and Tipton, 2014). In this article, we describe the introduction of a new reference vocabulary for the annotation of enzymes in UniProtKB – the Rhea knowledgebase of biochemical reactions (https://www.rhea-db.org) (Lombardot, et al., 2019; Morgat, et al., 2017). Rhea is a comprehensive expert-curated knowledgebase that uses the chemical ontology ChEBI (Chemical Entities of Biological Interest, https://www.ebi.ac.uk/chebi/) (Hastings, et al., 2016) to describe reaction participants, their chemical structures, and chemical transformations. Rhea provides stable unique identifiers and computationally tractable descriptions for around 12,000 unique biochemical reactions, including reactions of the IUBMB enzyme classification, as well as thousands of additional enzymatic, transport reactions, and spontaneous reactions. The introduction of Rhea as the reference vocabulary for enzyme annotation in UniProtKB will significantly improve the coverage and precision of enzyme annotation and will allow users of UniProtKB to leverage chemical structure data and classifications to answer biological questions.

In the following we describe the re-annotation of legacy enzyme data in UniProtKB using Rhea and a selection of the new tools and services we have developed to exploit this enhanced enzyme dataset.

## 2 Methods

### 2.1 Rhea as a reference vocabulary for enzyme annotation in UniProtKB

Prior to this work, UniProtKB used the hierarchical enzyme classification of the Enzyme Nomenclature Committee of the IUBMB (hereafter referred to simply as the enzyme classification) as the main reference vocabulary for enzyme annotation. The enzyme classification uses a hierarchy of exactly four levels to classify enzymes according to the chemistry of representative reactions, which are described in text form (Bairoch, 2000). In this work we introduce Rhea reaction identifiers as the reference vocabulary for enzyme annotation in UniProtKB, with the corresponding four digit enzyme class (EC number) selected from a distinct mapping of Rhea reactions to EC numbers (maintained by Rhea at https://www.rhea-db.org/download). The EC number annotations for Rhea reactions are optional in UniProtKB, as Rhea contains thousands of reactions not described by the IUBMB Enzyme Classification. Note that UniProtKB will continue to describe some enzymatic reactions in text form, particularly those reactions whose specific chemistry is not yet well characterized and which cannot be described using ChEBI.

### 2.2 Migration of legacy enzyme annotation in UniProtKB to Rhea and re-annotation

In order to lay the groundwork for the integration of Rhea in UniProtKB, we first mapped legacy textual descriptions of enzymatic reactions in UniProtKB to Rhea reaction identifiers. We accomplished this using the ENZYME database (Bairoch, 2000), which links textual reaction descriptions (as used in UniProtKB) to their corresponding EC numbers, and the Rhea database, which links EC numbers to their corresponding Rhea reactions. We checked and validated all such mappings of UniProtKB enzyme annotation – EC number – Rhea identifier derived in this way. A small number of legacy UniProtKB enzyme annotations were not based on EC numbers, and we mapped these textual descriptions manually to Rhea identifiers where possible, creating new Rhea reactions where needed. We then used the completed mapping to replace the legacy textual descriptions of enzymatic reactions in UniProtKB by Rhea annotations, and to update all automatic annotation rules that are used to add enzyme annotations to UniProtKB/TrEMBL records, including those from HAMAP (Pedruzzi, et al., 2015) and PROSITE (Sigrist, et al., 2013). Mapping of all EC number annotations is now complete, while the mapping of additional legacy enzyme data described in natural language in UniProtKB/Swiss-Prot (mainly in “FUNCTION” annotations) is still ongoing.

## 2.3 UniProt tools and services that use Rhea

We modified the UniProt data model and output formats – text, XML, and RDF – to include Rhea reaction data and references to ChEBI. We modified the UniProt website https://www.uniprot.org/ (Jain, et al., 2009), UniProt REST API https://www.uniprot.org/help/api and SPARQL endpoint https://sparql.uniprot.org/ to support searches using Rhea and ChEBI identifiers as well as ChEBI names, synonyms, and chemical structures represented as InChIKeys. The InChIKey, a simple hash representation of chemical structures, provides a convenient means to search and map chemical structure databases. The first layer of the InChIKey encodes information on connectivity, the second layer encodes information on stereochemistry, and the third encodes information on charge. A more complete description of the InChIKey is available at https://www.inchi-trust.org/.

## 3 Results

### 3.1 Annotation of UniProtKB using Rhea

We performed a complete re-annotation of legacy UniProtKB enzyme data using Rhea (as described in Methods), and now use Rhea as the primary reference vocabulary for enzyme annotation in UniProtKB. UniProtKB/Swiss-Prot currently includes annotations for 5,795 unique Rhea reactions (around 49% of all Rhea reactions), which feature in 215,789 distinct UniProtKB/Swiss-Prot protein records (38.5% of all UniProtKB/Swiss-Prot records are annotated with Rhea) (release 2019_05 of June 5 2019). Of the 5,795 Rhea reactions used in UniProtKB/Swiss-Prot, 4,828 reactions (around 85% of all reactions) are linked to EC numbers. We are currently working to improve the coverage of the 6,140 Rhea reactions not currently represented in UniProtKB/Swiss-Prot through a variety of approaches. These approaches include continuing expert curation of new literature, re-annotation of free text legacy annotations in UniProtKB/Swiss-Prot entries, and integration of data from other resources that use Rhea. One such resource of note is the SwissLipids knowledgebase (Aimo, et al., 2015), which contains annotations for more than 2,000 unique Rhea reactions that are not yet represented in UniProtKB. We will describe these and other annotation efforts in more detail in forthcoming publications.

### 3.2 UniProt tools and services that use Rhea

Below we describe how users can navigate and exploit Rhea data using the UniProt website, REST API, and SPARQL endpoint.

#### 3.2.1 Rhea and the UniProt website

The UniProt website https://www.uniprot.org constitutes the main point of entry for most UniProt users. The UniProtKB entry view provides a summary of annotated Rhea reactions for each enzyme (Figure 1). Users can choose to reveal the two-dimensional structures of reaction participants for each annotated reaction, and click on the reactions and their participants to launch searches in UniProtKB or Rhea or link out to ChEBI.

**Figure 1.**
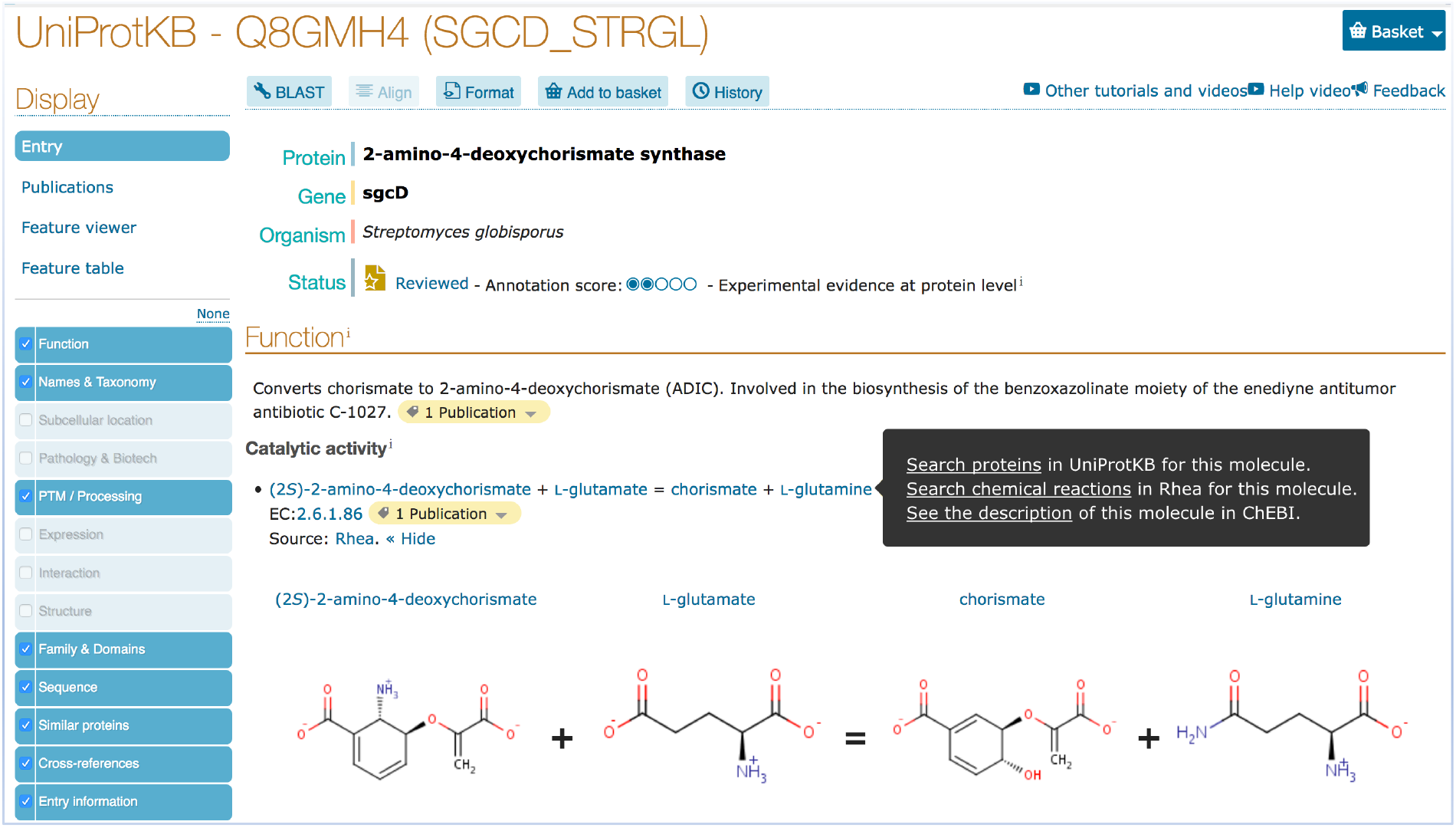
UniProtKB entry view showing Rhea annotation for the Streptomyces globisporus enzyme 2-amino-4-deoxychorismate synthase (UniProt:Q8GMH4 annotated with RHEA:25512). The search and link-out options available for each reaction participant are illustrated using “L-glutamine”; users can search in UniProtKB or Rhea, or link out to ChEBI to learn more about the metabolite in question. We omit most sections for clarity.

Figure 2 illustrates selected advanced search options for reactions, chemical names and structures in UniProtKB. Users can search for identifiers from Rhea as well as identifiers, names, synonyms (Figure 2a), and chemical structures (encoded as InChIKeys) from ChEBI (Figure 2b). The complete ChEBI ontology is indexed to support hierarchical searches, while ChEBI identifiers entered by users are mapped to those of the major species at pH 7.3, the form used in Rhea, using the mapping provided at https://www.rhea-db.org/download. InChIKeys provide a means to query UniProtKB using chemical structure data from other resources, including reference knowledgebases such as the Human Metabolome Database (HMDB) (Wishart, et al., 2018) or LIPID MAPS (Fahy, et al., 2009). Users of these and other resources can simply convert their structures to InChIKeys and use them to query UniProtKB. Searches may be performed using the complete InChIKey (to find exact structure matches), or using the first and second layers of the InChIKey (to find molecules with matching connectivity and stereochemical orientation, irrespective of charge, as in Figure 2b), or using only the first layer of the InChIKey (to find molecules with matching connectivity).

**Figure 2.**
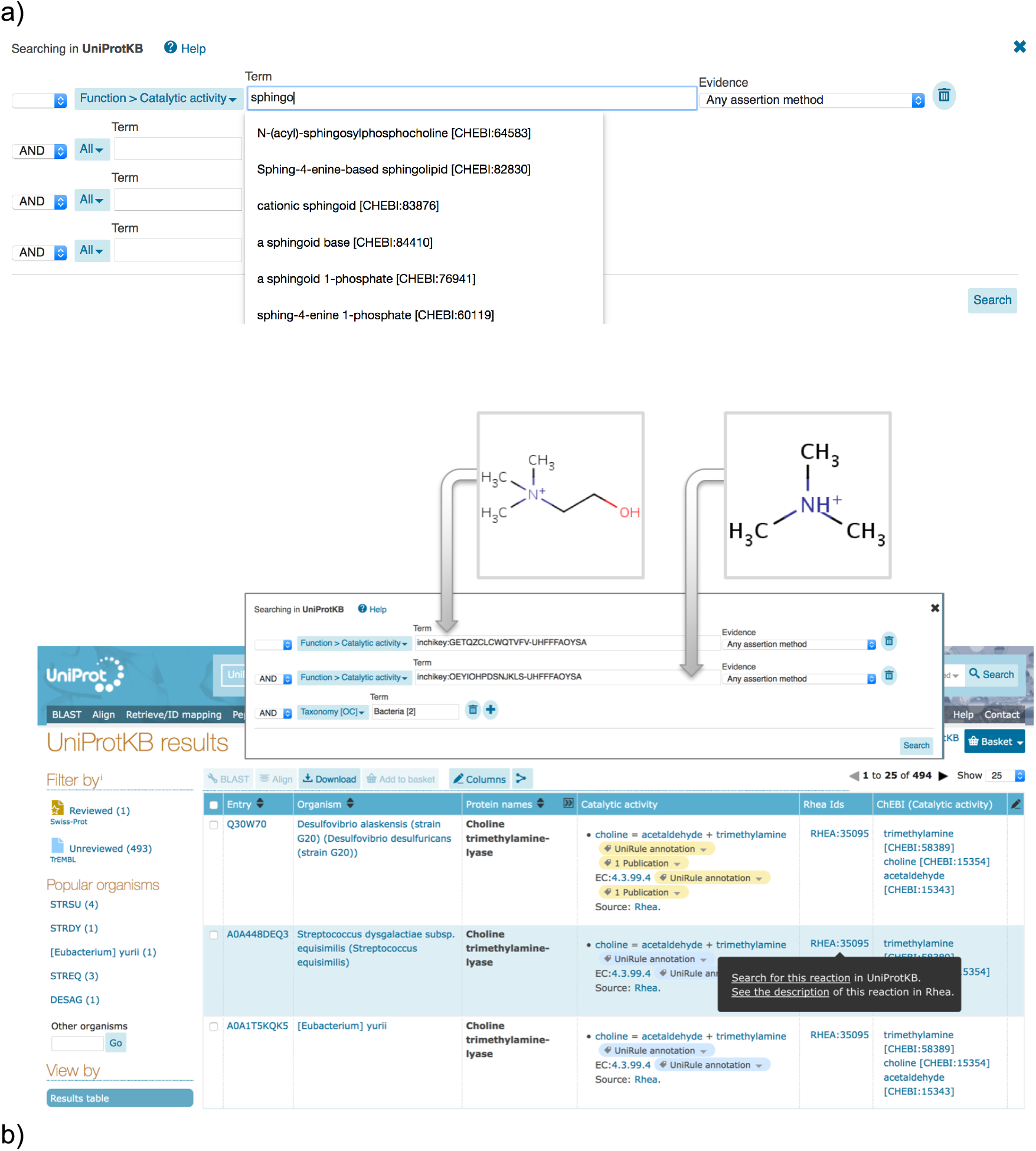
Advanced search in UniProtKB for enzymatic reactions. a) Advanced search using chemical nomenclature. The autocomplete feature is shown. b) Advanced search using InChIKeys of choline (HMDB00097) and trimethylamine (TMA) (HMDB0000906) to identify bacterial enzymes metabolizing both compounds. The result table can be customized to display Rhea reaction data, which can be used to launch further searches and link out precisely as in the entry view.

Figure 2b illustrates an InChIKey search for microbial enzymes that metabolize choline (HMDB00097, InChIKey=OEYIOHPDSNJKLS-UHFFFAOYSA-N) and trimethylamine (TMA) (HMDB0000906, InChIKey=GETQZCLCWQTVFV-UHFFFAOYSA-N). Host gut microbes produce TMA from choline, while the human liver enzyme FMO3 then converts TMA to the pro-atherogenic molecule trimethylamine N-oxide (TMAO) (HMDB0000925) (Canyelles, et al., 2018; Chhibber-Goel, et al., 2017). This InChIKey search allows users of HMDB to exploit protein annotation in UniProtKB to connect the bacterial choline trimethylamine-lyase *cutC* of *Desulfovibrio alaskensis* (UniProtKB:Q30W70) and homologs to human enzymes such as FMO3 by their common metabolites.

For those chemical structures that have no corresponding match annotated in UniProtKB – no matching InChIKey first layer – users might choose instead to search for relevant information about the chemical classes to which these structures belong. Chemical structures can be mapped to their corresponding ChEBI classes using tools such as ClassyFire (Djoumbou Feunang, et al., 2016), and the ChEBI classes then used to search UniProtKB. Interested readers can find further information about chemical data search in UniProtKB in the online documentation provided at https://www.uniprot.org/help/chemical_data_search.

#### 3.2.2 Rhea and the UniProt REST API

The UniProt website serves a REST API (https://www.uniprot.org/help/api) that allows users to query and process data programmatically. The REST API has been modified to handle Rhea and ChEBI identifiers, as well as ChEBI names, synonyms, and structural data. The sample query below will retrieve the aforementioned bacterial enzymes metabolizing choline and trimethylamine using their respective InChIKeys (see also https://tinyurl.com/y2mcjotd).

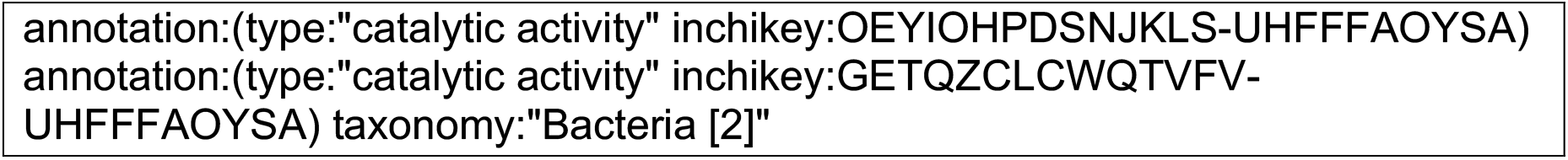

As before, this particular query uses only the first and second layers of the InChIKey to allow permissive matching between different charge state representations. Users can modify REST queries like that shown above in order to specify both the required data output (annotation fields) and format (such as. tab,. xls,. rdf, and others) (for more details see https://www.uniprot.org/help/api_queries).

#### 3.2.3 Rhea and the UniProt SPARQL endpoint

The <u>UniProt SPARQL endpoint</u> allows users to perform complex federated queries that combine RDF data from UniProt with that from other SPARQL endpoints. Like the UniProt website and REST API, the UniProt SPARQL endpoint now supports queries using Rhea and ChEBI identifiers, as well as ChEBI names, synonyms, and structural data. Figure 3 provides a sample SPARQL query that combines the UniProt, Rhea (Lombardot, et al., 2019) and ChEMBL (Gaulton, et al., 2017) SPARQL endpoints. The aforementioned pro-atherogenic metabolite TMAO, produced from choline and TMA by a combination of microbial and human enzymes, may promote atherosclerosis by perturbing enterohepatic cholesterol metabolism and macrophage cholesterol transport (Canyelles, et al., 2018; Chhibber-Goel, et al., 2017). This query will find drugs in ChEMBL that target human enzymes acting on sterols (members of the ChEBI class CHEBI:15889).

**Figure 3.**
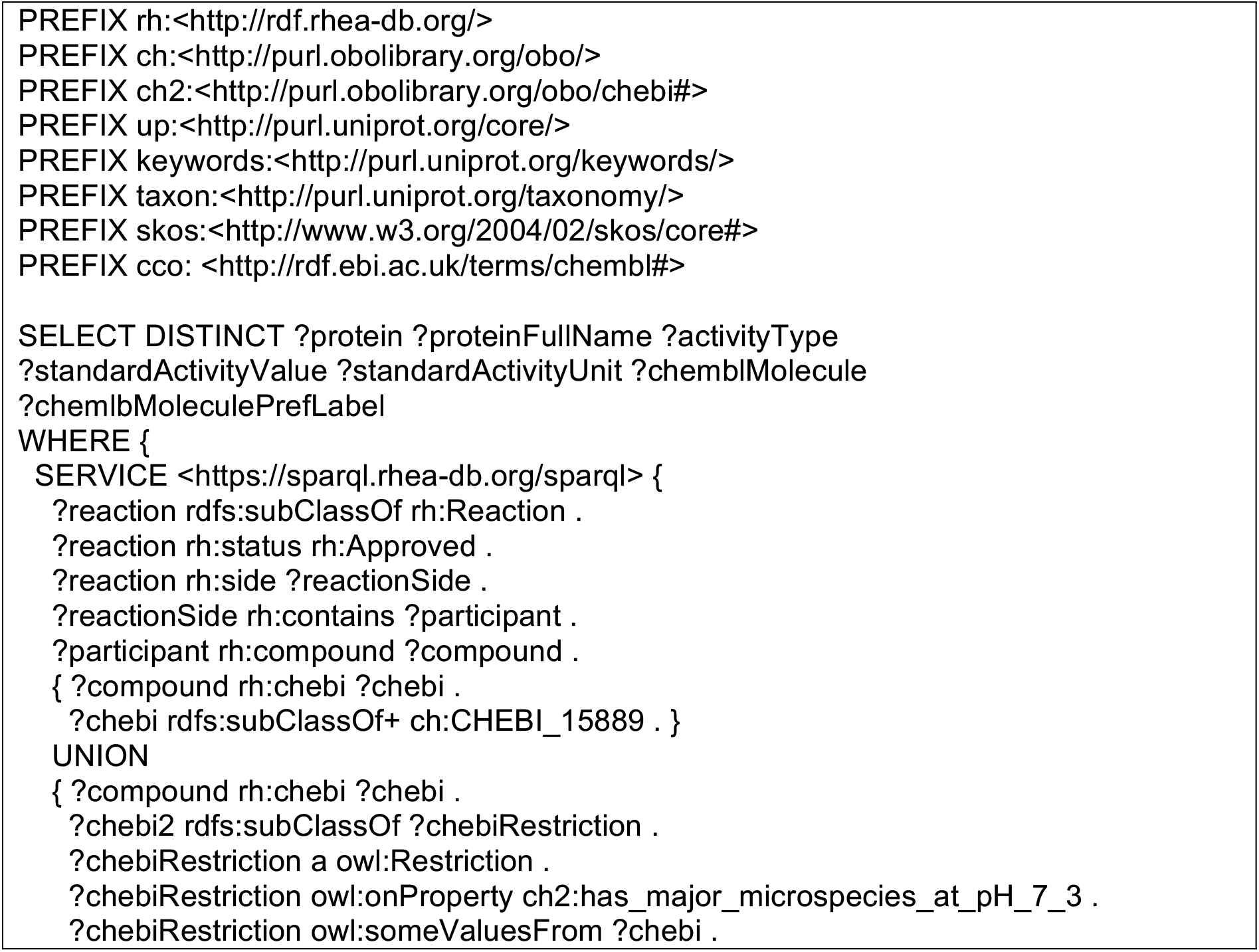
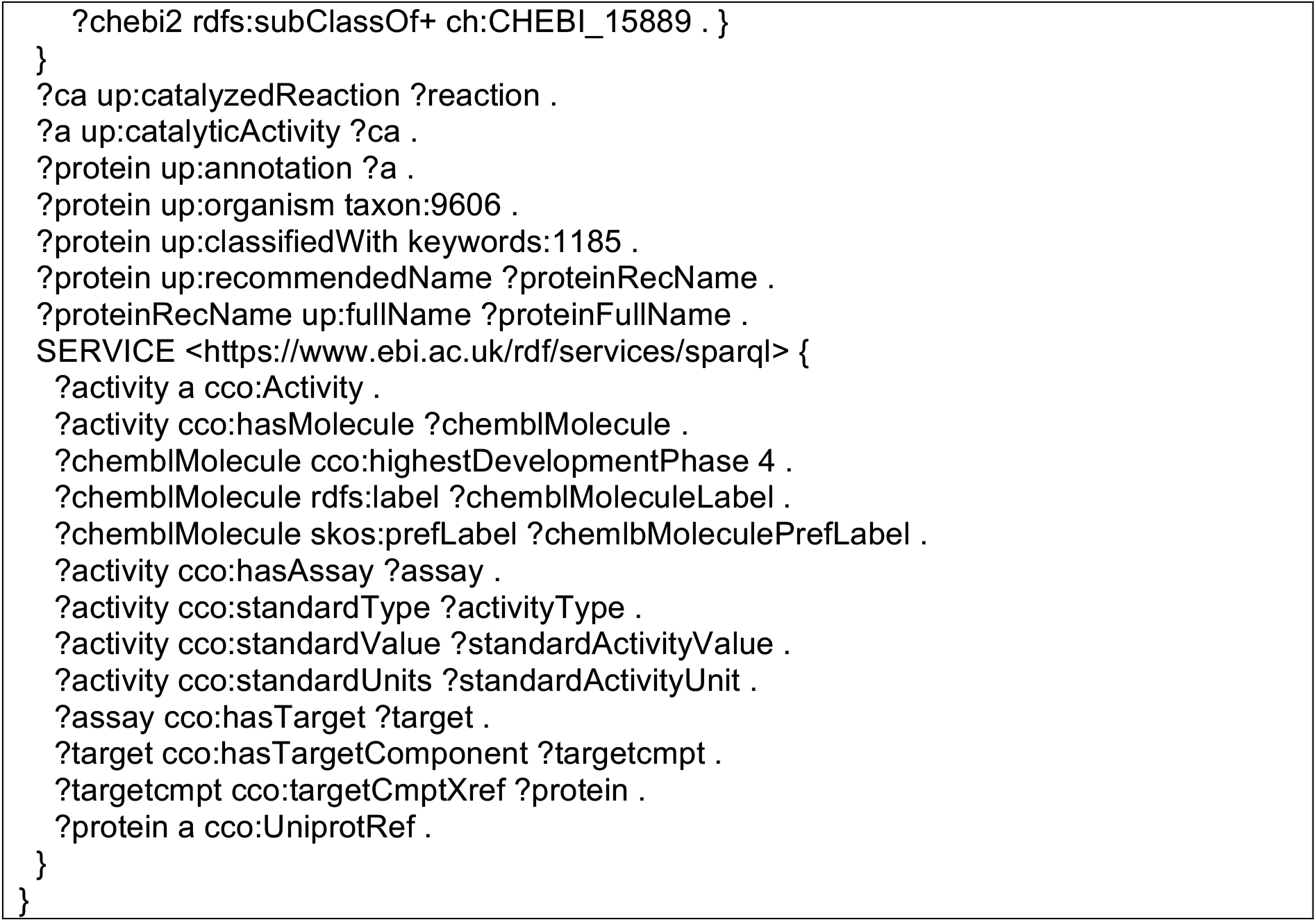
A sample federated SPARQL query that leverages Rhea annotation in UniProtKB. The query retrieves information about drugs that target enzymes involved in human sterol (CHEBI:15889) metabolism from the UniProt, Rhea and ChEMBL SPARQL endpoints, federating the three SPARQL endpoints with two SERVICE calls.

Other federated queries might explore metabolism in the context of genomic organization and variation using resources such as Ensembl (Zerbino, et al., 2018) or how metabolism evolves using resources such as OMA (Altenhoff, et al., 2018) and OrthoDB (Kriventseva, et al., 2019). We describe more sample queries in the documentation available at the UniProt SPARQL endpoint.

## 4 Conclusions and futures directions

Here we describe the introduction of Rhea as the reference vocabulary for enzyme annotation in UniProtKB. This development will significantly increase the utility of UniProtKB for applications such as the integrative analysis of metabolomics and other ‘omics data (Kale, et al., 2016; Sud, et al., 2016), as illustrated here, as well as the study of enzyme chemistry and evolution (Rahman, et al., 2016; Tyzack, et al., 2019), the construction and annotation of metabolic models (Cottret, et al., 2018; King, et al., 2016; Moretti, et al., 2016), and the engineering of pathways for biosynthesis and bioremediation (Duigou, et al., 2019). Rhea will also provide a basis to further improve links with other knowledge resources, such as the Gene Ontology (The Gene Ontology Consortium, 2019) and Reactome (Fabregat, et al., 2018), which both plan to use Rhea as the reference for reaction chemistry (Chris Mungall and Peter D’Eustachio, personal communication). We also plan to extend the use of ChEBI within UniProtKB to describe all small molecule chemical structure data, including functionally important ligands and post-translational modifications (see https://www.uniprot.org/docs/ptmlist). This will further improve links and interoperability between UniProtKB and other resources that use ChEBI such as the metabolomics data repository MetaboLights (Kale, et al., 2016), the IMEx molecular interaction databases (Orchard, et al., 2012) and the Complex Portal (Meldal, et al., 2015), the literature annotation services of Europe PubMed Central (Europe PMC Consortium, 2015), the BioModels repository (Glont, et al., 2018), and the Immune Epitope Database (IEDB) (Dhanda, et al., 2019).

## Acknowledgements

We thank Marco Pagni of the Vital-IT group of the SIB Swiss Institute of Bioinformatics for critical reading of the manuscript and helpful comments, and the Cheminformatics and Metabolism Team of EMBL-EBI for their work in maintaining and developing ChEBI.

